# Sexual selection in capelin: endurance rivalry insights from semen quality and regeneration

**DOI:** 10.1101/2024.06.06.597742

**Authors:** Ranjan Wagle, Connor P. Hanley, Nick C. Murphy, Craig F. Purchase

## Abstract

Male-biased adult sexual size dimorphism is often the result of intra-sexual selection driven by male-male competition. Capelin (*Mallotus villosus*) exhibit male-biased sexual size dimorphism but lack contest competition or clear scramble competition, and female mate choice, if present, is unrelated to male size. Consequently, it is hypothesized that adult sexual size dimorphism in capelin is due to a mating system that favours larger males with greater energy reserves, allowing them to compete for prolonged access to mating opportunities through endurance rivalry. To enable this, males need a continual supply of semen throughout the spawning season. However, they have unusually small testes and are predicted to deplete stored semen rapidly. Furthermore, their unique sperm physiology (i.e. the only known externally fertilizing vertebrate to release pre-activated semen) may constrain their ability to regenerate it. We found that the majority of capelin sampled on beaches had adequate semen, but towards the end of the spawning season, some fish had none. Capelin held in laboratory tanks could regenerate semen within two days, maintaining their ability to seize continual mating opportunities. These results support the endurance rivalry hypothesis.

## Introduction

Sexual selection “arises from fitness differences associated with nonrandom success in the competition for access to gametes for fertilization” (Shuker and Kvarnemo 2021), and occurs both before and after mating. It is based on intra-sex competition and excludes forms of “narrow sense” natural selection such as ‘viability selection’ (survival) and ‘fecundity selection’ (the ability to produce gametes). Pre-mating mechanisms include intra-sex contest competition, scramble competition and endurance rivalry, and inter-sex mediated intra-sex competition through mate choice (Shuker and Kvarnemo 2021; Purchase et al. 2026). These act asymmetrically on males and females and can lead to differences in their phenotypes, including body size.

Sexual size dimorphism (SSD) refers to adult body size differences between males and females within a species and reflects sex-specific selection pressures on body size. SSD is widespread across the animal kingdom, and its magnitude and direction vary among taxa depending on the relative natural and sexual selection pressures on each sex (Darwin 1872; Andersson 1994; Rennie et al. 2008; Horne et al. 2020). Female-biased SSD (adult females being larger than males) is commonly attributed to natural selection (‘fecundity selection’), favouring large females in egg production capacity (but see Shine 1988). In contrast, male-biased SSD is often the result of sexual selection driven by male-male competition (Trivers 1972; Fairbairn 1997; Cox et al. 2003; Pyron et al. 2013). The majority of fish species exhibit female-biased SSD (Bisazza 1993; Webb and Freckleton 2007), but there are exceptions that can provide insights into selection dynamics and mating systems (Parker 1992; Bisazza 1993; Brockmann 2001; Orbach et al. 2019).

Capelin (*Mallotus villosus*) is one such atypical species (Templeman 1948; Orbach et al. 2019). In addition to being 10-20% larger than females of the same age, adult males also exhibit secondary sexual characteristics, such as relatively enlarged fins and unique spawning ridges (Pitt 1958; Winters 1970; Jangaard 1974; DFO 1991; Orbach et al. 2019). These have recently been hypothesized to be under natural selection by facilitating physical contact with females while spawning, rather than inter-sex mediated sexual selection through mate choice (Orbach et al. 2019). Despite the male-biased SSD in capelin evidence does not support contests/fighting for mating opportunities, and if females do exhibit some sort of mate choice, it is not based on male size (Orbach et al. 2020). It is likely that large males benefit in some way, as achieving a larger size usually requires longer periods of growth or increased foraging risk (Rennie et al. 2008); thus, there is a trade-off between the benefits of size and the survival probability costs associated with delaying reproduction (Roff 1986).

Scramble competition is framed as leading to traits that enable males to more quickly locate mates (Andersson 1994; Dechaume-Moncharmont et al. 2016). This does not occur in capelin as both sexes arrive at the same time in huge abundances at spawning beaches. Capelin spawn on both beach (inter-tidal zone) and off beach (benthic) sites. Beach-spawning capelin are easily observed to spawn as couples or threesomes (Templeman 1948; DFO 1991; Orbach et al. 2019), however when off the beach it is difficult to observe what happens, as very large groups of fish are together. Males may ‘scramble’ on top of females similar to garter snake (*Thamnophis sirtalis parietalis*) mating balls (Shine et al. 2000, 2003). Larger body size and larger relative fin size may enable greater maneuverability, a potential sexual selection mechanism not previously considered (Orbach et al. 2019, 2020).

Orbach et al. (2019) proposed that capelin male-biased SSD is due to a mating system based on endurance rivalry, where male-male competition is interaction-independent (Andersson 1994), and the ability to be present at spawning sites for longer is adaptive (e.g., European treefrog *Hyla intermedia*; Castellano et al. 2009). Capelin migrate inshore to spawn, and once it begins males remain in close proximity to the beach as spawning occurs intermittently with changing environmental conditions and the arrival of new females (Jeffers 1931; DFO 1991). By staying at the spawning site for longer periods, males would increase their chances of mating, which may be extended by larger bodies having more energy reserves. However, simply remaining at the site is insufficient as they also require enough semen to secure multiple matings. This could be achieved by possessing a large amount of semen at the onset of the spawning season or by producing new semen for each mating. However, capelin have unusually small testes with a gonadosomatic index of around 1% (Beirão et al. 2015; Orbach et al. 2020), and thus, the former does not occur. Capelin also possess unique semen, being the only known externally fertilizing vertebrate to release pre-activated sperm (Beirão et al. 2018). This may constrain their ability to regeneration sperm. If the mating system is based on endurance rivalry, once spawning begins at a site we predicted that (1) if sampled next to spawning beaches, some males will not contain any semen (it gets used up), but (2) they can quickly regenerate it despite their unique sperm physiology.

## Materials and methods

### Among-male variation in semen quality – spawning beach surveys

Spawning male capelin were captured during June and July 2023, using a dip net on three beaches of the Avalon Peninsula, Newfoundland, Canada: Middle Cove Beach (47° 39’ 2.75" N, 52° 41’ 45.24" W) on 29^th^ June; 2^nd^, 4^th^, 6^th^,10^th^, 12^th^ and 14^th^ July; Bellevue Beach (47° 38’ 9.28" N, 53° 46’ 51.09" W) on 7^th^, 8^th^, 9^th^ July; and Holyrood Beach (47° 23’ 21.8" N, 53° 07’ 34.5" W) on 21^st^ July. Fish from these beaches are thought to be from the same population (Dodson et al. 1991; Penton et al. 2014).

At capture, the total length of haphazardly selected males was measured, and semen was assessed by gently pressing the abdomen until no further release of semen occurred. Semen quality was visually scored immediately following release on to the fish’s body in the field according to the categories: excellent (white/dense semen – relatively high sperm concentration), fair (watery/dilute semen – relatively low sperm concentration), poor (thick-pelleted semen), or none (no semen). Although untested, we believe the poor quality semen category does not have the ability to fertilize eggs or be released during ejaculation. For excellent and fair semen qualities, the quantity was scored as relatively high, medium, or low. This categorization of semen quality and quantity was based on many years of observations with thousands of capelin and represented a compromise between resolution and repeatability.

To confirm the effectiveness of the semen quality categorization, four males were selected: two with excellent and two with fair semen quality. Using equal amounts of semen, excellent quality males had, on average, 6.75 times higher fertilization success than those deemed of fair quality, when bred with the same females (see Wagle 2024).

### Semen regenerative capacity – lab experiment with wild fish

Semen regenerative capacity (the ability to replenish semen stock) was assessed in a blocked design comprised of four blocks of capelin captured at Middle Cove on June 29, July 4, and July 10, and Holyrood Beach on July 21. Fish were transported in aerated coolers to the Ocean Sciences Centre (OSC), Memorial University of Newfoundland, St. John’s, NL, Canada.

Within each block, all males (∼ 80) were measured for total length and completely stripped of their semen (day 0), a subset of which was then scored for quality and quantity using the categories previously described. Each block was held in a separate 1000L flow-through seawater tank (four tanks total), maintained at 4 to 8 °C. Over a period of six days (total duration of captivity), 10-20 males were selected randomly on alternating days and stripped again (days 2, 4 and 6 post-capture) to score regenerative semen quality and quantity following the same assessment categories and then discarded. Fish tagging was not required, as each male was stripped and assessed exactly twice (day 0 and once later) before discard. The first block (Middle Cove, June 29) deviated from this alternate-day pattern, as assessments were conducted on days 2, 3, and 4. No food was provided, as capelin generally do not feed while spawning. No individuals died during captivity.

### Data analysis

Graphs depicting the patterns of semen quality, quantity and regenerative capacity during the spawning period were plotted using R v. 4.3.0 (R Core Team 2023) to assess variations in semen quality and quantity. Data were adjusted by converting to proportionate scores (number of fish in each semen quality category divided by total number of fish scored in each date and location) for consistent comparisons across semen quality and quantity, capture date and location. Box plots were used to depict the relationship between capelin total length, semen quality and quantity, and semen regenerative capacity. We report descriptive summary statistics rather than inferential tests, as the latter are inappropriate due to small the sample size and confounding spatiotemporal variation.

## Results

### Among-male variation in semen quality

Across 11 sampled dates (June 29 to July 21), semen quality showed notable inter-individual variation (Fig. 1). The majority of sampled capelin (average 53%) had excellent quality semen, ranging from a maximum of 90% on July 4 to a minimum of 34% on July 14 (Fig. 1). Fair quality semen was found in, on average, 31% of the fish, reaching a maximum of 56% on July 2 and a minimum of 8% on July 10. Only 9% of sampled capelin had poor-quality semen. About 7% of sampled capelin had no semen, which was not observed until July 9.

**Fig. 1.**
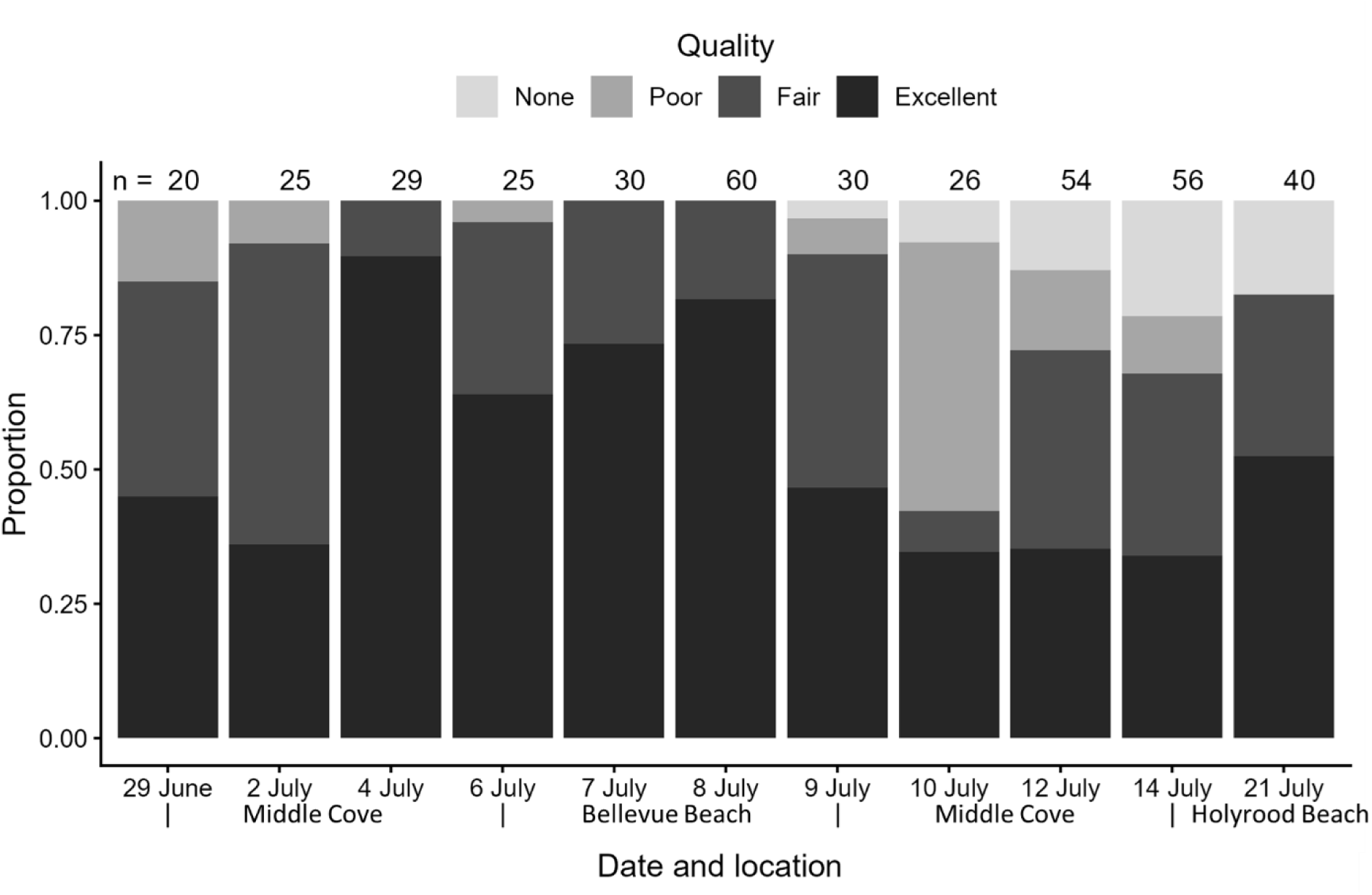
Variation in capelin semen quality scored in the field at different beaches in eastern Newfoundland during the 2023 spawning period. Semen quality was categorized into four categories, each denoted by different shades. The total number of capelin scored on specific dates is indicated above each respective bar. The y-axis represents proportionate scores, standardized through conversion to proportionate values, normalizing the data for comparison.

Around 16% of the sampled males that had poor quality semen or no semen (Fig. 1) are believed to have lacked the ability to fertilize any eggs at the time of sampling. The proportions of excellent and fair-quality semen fluctuated without a clear temporal trend, while instances of capelin with no semen increased slightly over the spawning period. However, interpreting a temporal trend is challenging due to the small sample size and confounding factors of time and space.

Capelin with excellent semen quality were often accompanied by relatively high quantities (76%; Fig. 2), whereas 54% had a medium quantity of fair quality semen. Very few capelin with relatively low quantity semen were present across both quality categories (3% in excellent while 9% in fair quality semen).

**Fig. 2.**
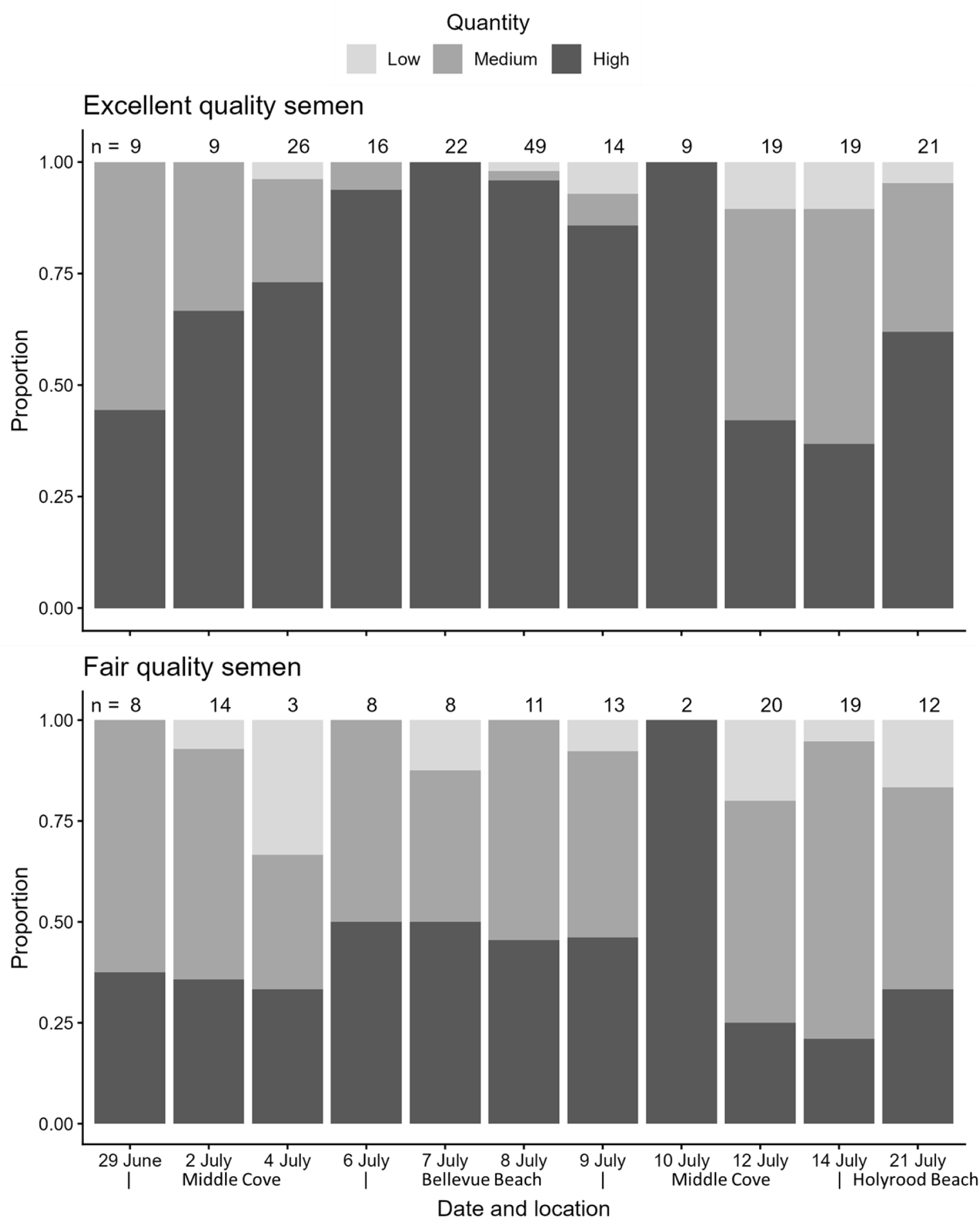
Variation in capelin semen quantity within the excellent and fair quality categories scored in the field across different beaches in eastern Newfoundland during the 2023 spawning period. Relative semen quantity was classified into three categories, each denoted by different shades. The total number of capelin scored within each of the two quality categories on specific dates is indicated above each bar. The y-axis represents proportionate scores, standardized through conversion to proportionate values, normalizing the data for comparison.

There was no relationship between semen quality and male size (Fig. 3). Body size was not related to the quantity of fair quality semen, but larger males had higher quantities of excellent quality semen compared to smaller males (Fig. 4).

**Fig. 3.**
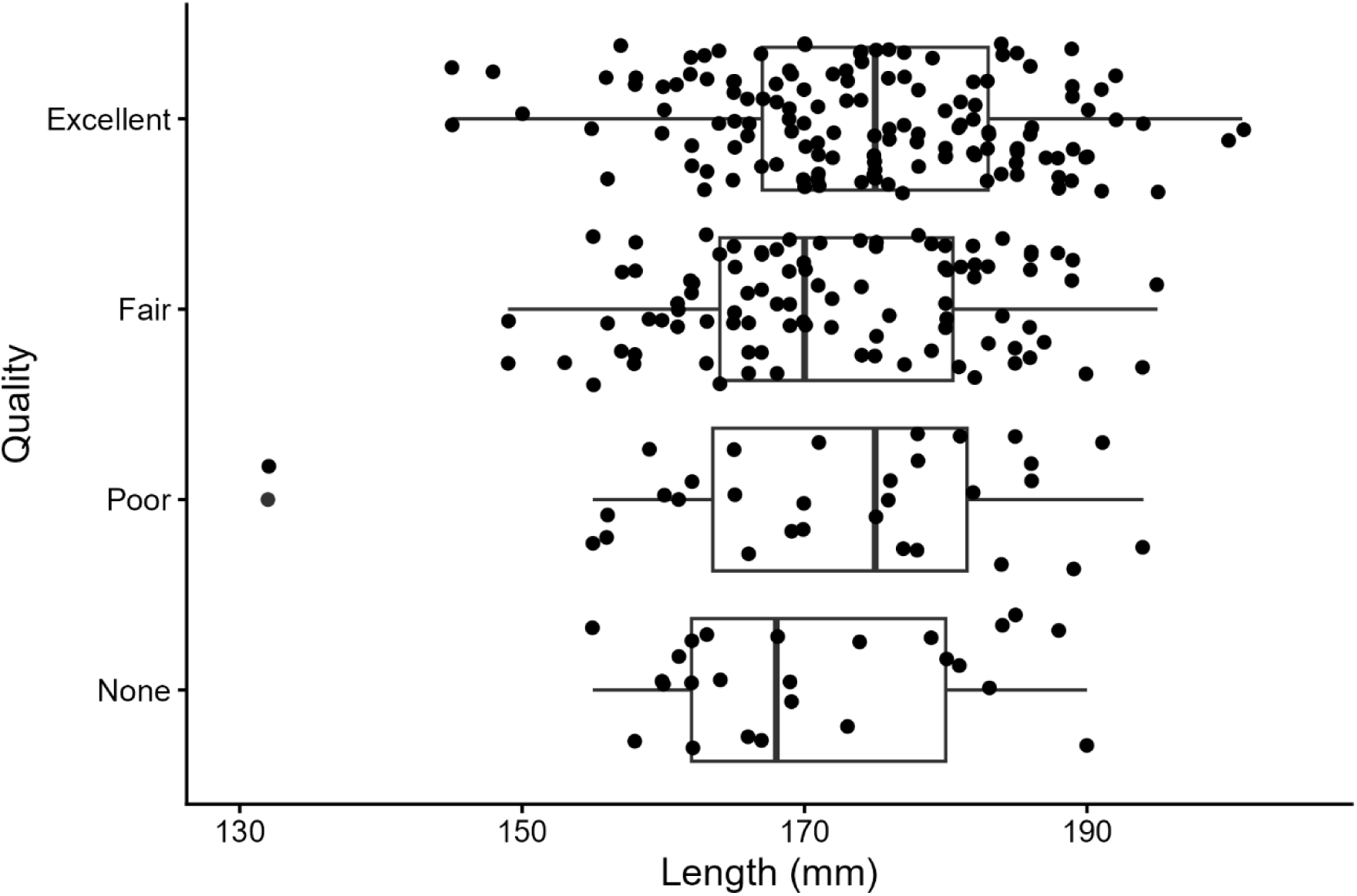
Variation in capelin semen quality across different fish total lengths scored in the field across different beaches in eastern Newfoundland during the 2023 spawning period. The thick line within the box plot represents the median, the box represents the interquartile range (25th and 75th quartiles), and the whiskers extend to 1.5 times the interquartile range from the first and third quartiles, respectively.

**Fig. 4.**
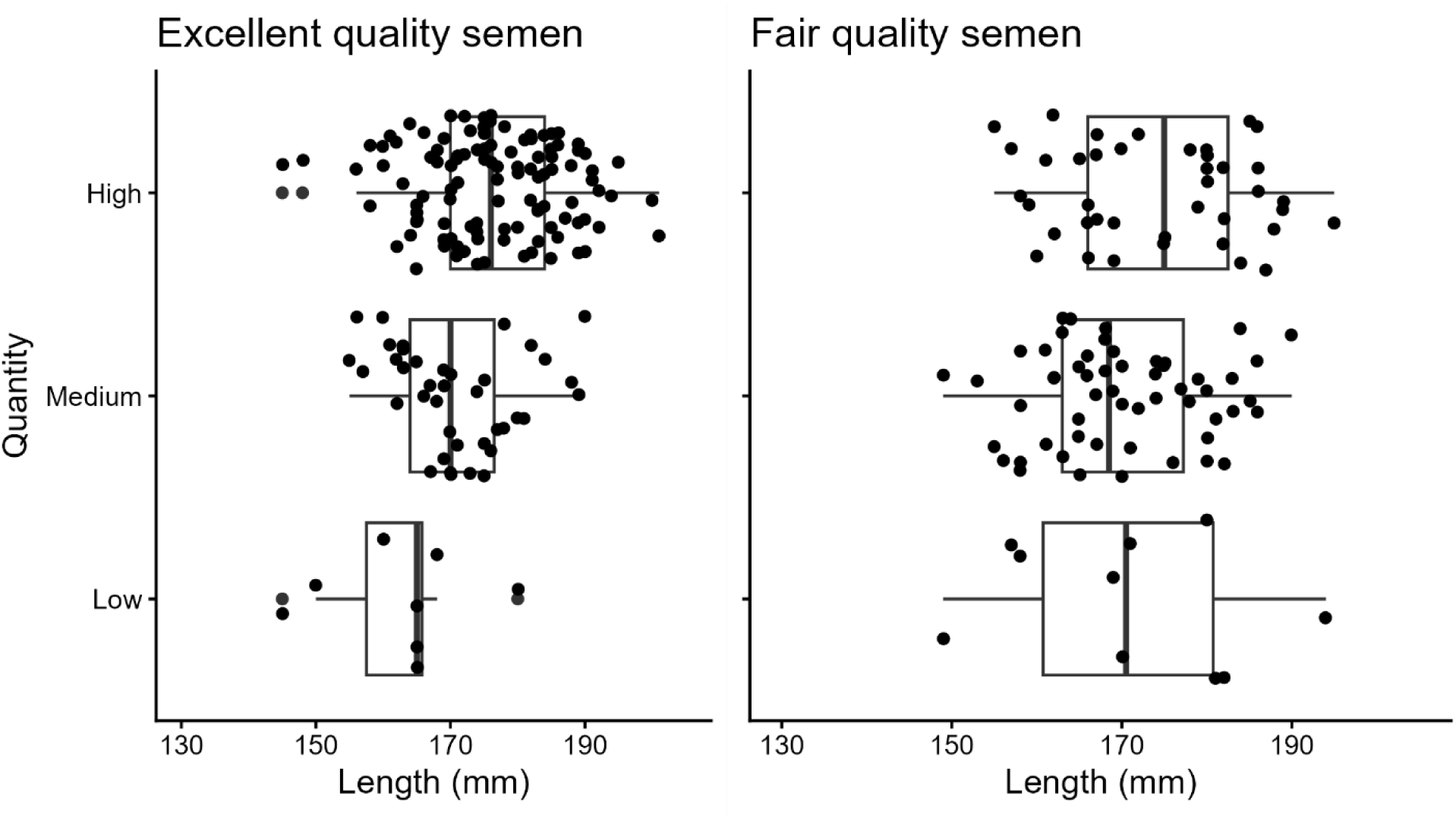
Variation in capelin semen quantity across different fish total lengths in two semen quality categories, excellent and fair, scored in the field across different beaches in eastern Newfoundland during the 2023 spawning period. The thick line within the box plot denotes the median, the box represents the interquartile range (25th and 75th quartiles), and the whiskers extend to 1.5 times the interquartile range from the first and third quartiles, respectively.

### Semen regenerative capacity

Across all four capture dates (blocks), the majority of assessed males (Fig. 5) had semen of excellent (average 53%) and fair (average 21%) quality at the time of capture (day 0). Many individuals were able to regenerate excellent (36%) and fair (41%) quality semen after two days (was not assessed on day 1) post-strip across all blocks. However, that proportion only increased (Fig. 5) in the next assessment in two blocks (June 29 and July 19, Middle Cove). Individuals captured on June 29 could regenerate excellent and fair quality semen the fastest amongst all four blocks. Although the fish were not tracked individually, it is likely that fish with poor quality semen upon capture also had poor quality on subsequent days. Individuals from July 4 could regenerate faster than those from July 10, however, those from July 10 initially contained lower semen quality than other blocks. Individuals from July 21 showed a reduced ability to regenerate excellent and fair quality semen over subsequent days.

**Fig. 5.**
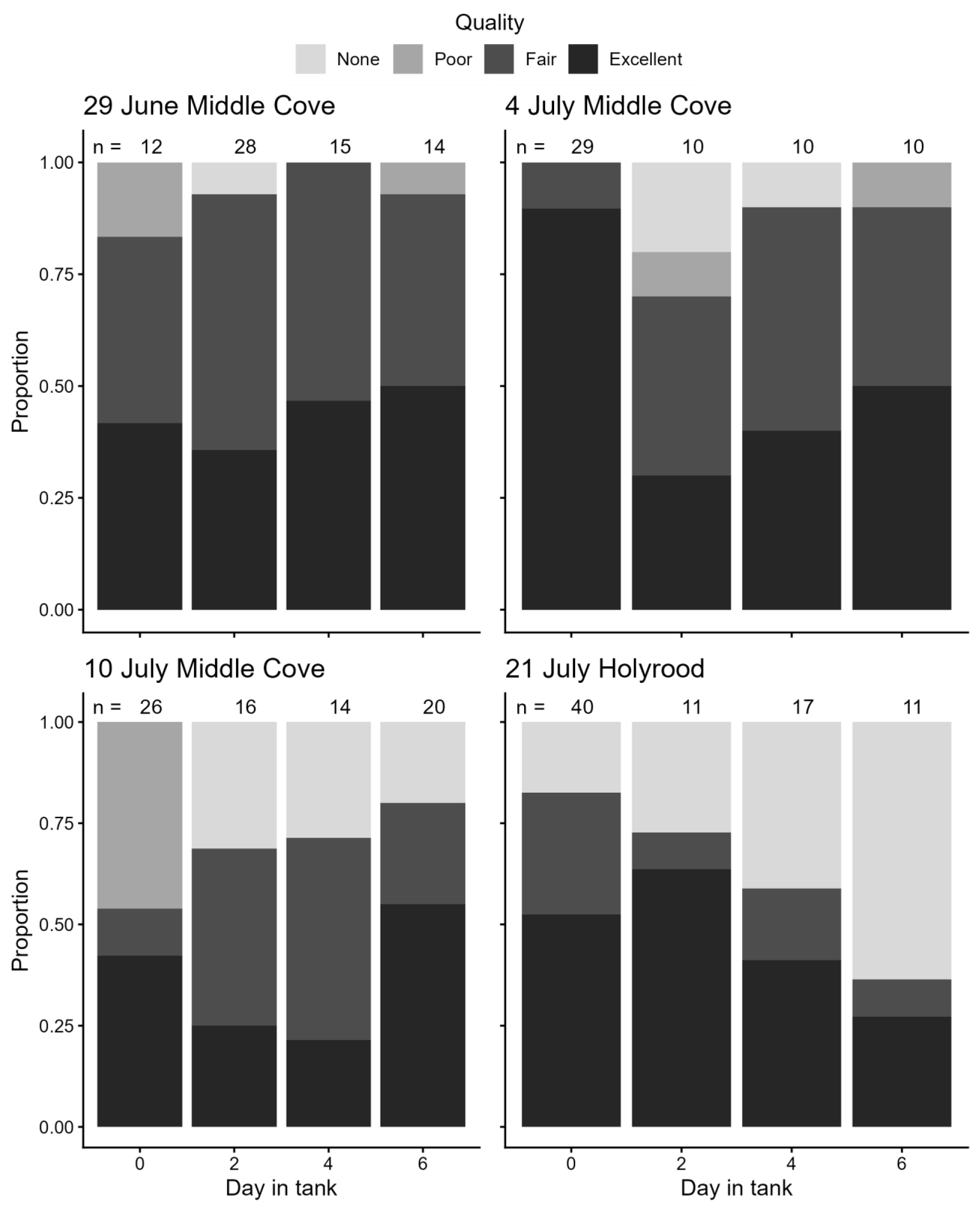
Assessment of the regenerative capacity of capelin semen based on quality. Completely stripped (Day 0) male capelin were kept in a tank, and every alternate day over a 6-day period, a subset was scored to assess their semen regeneration ability based on four categories of semen qualities, each distinguished by distinct shades. The total number of capelin scored on specific dates is indicated above each respective bar. The y-axis represents proportionate scores, standardized through conversion to proportionate values, normalizing the data for comparison. The capture date and location are provided at the top of each bar plot.

For excellent and fair quality semen (Fig. 6; Fig. 7), most individuals contained high and medium quantities of semen on day 0 (excellent quality: 68% high and 29% medium quantities; fair quality: 39% high and 48% medium quantities). Individuals captured on June 29 and July 4 that were able to regenerate excellent quality semen on subsequent days exhibited greater quantitative regeneration of semen compared to those of the same ability captured on July 10 and July 21 (Fig. 6). A similar pattern was observed in individuals that produced fair quality semen (Fig. 7).

**Fig. 6.**
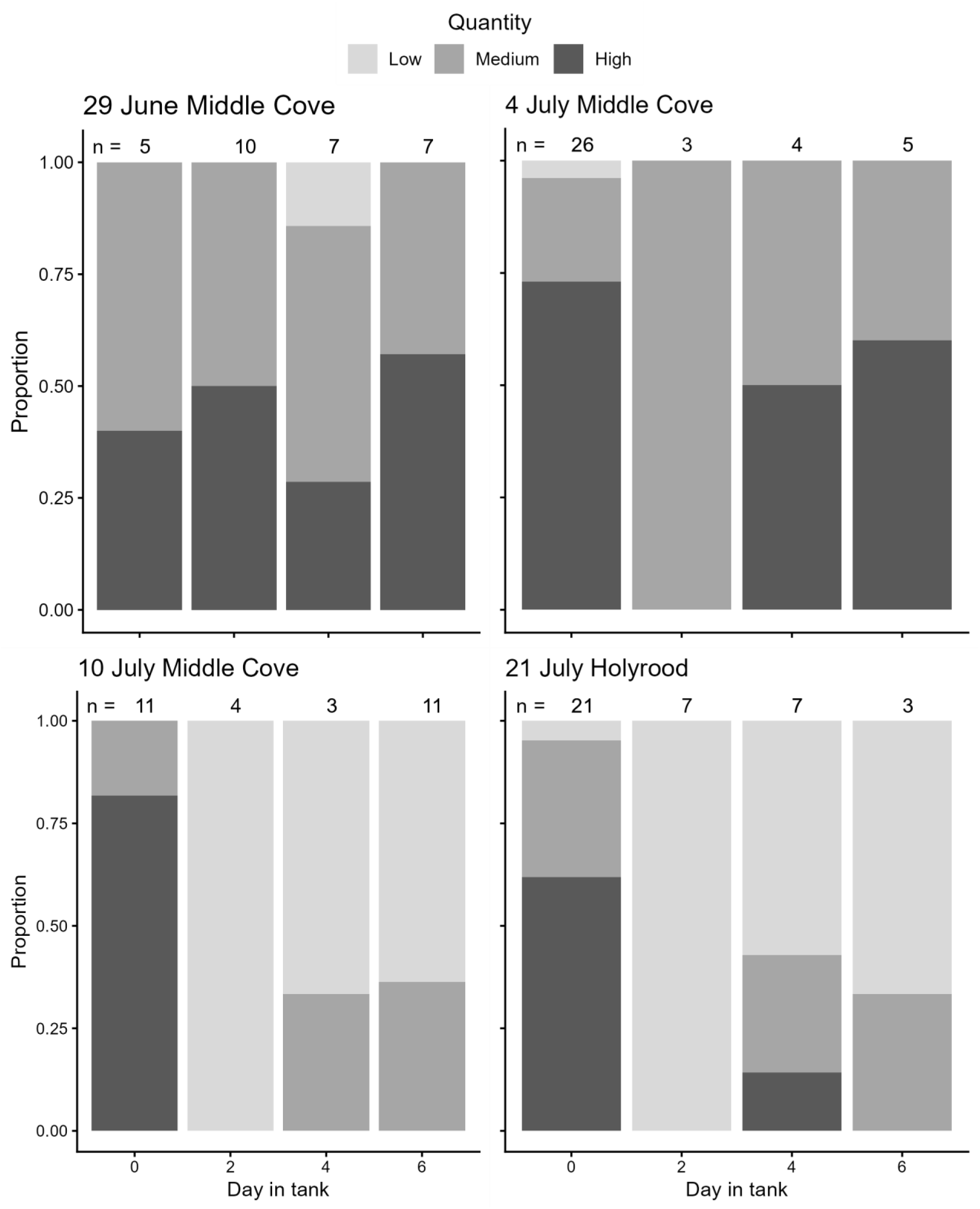
Assessment of the regenerative capacity of excellent quality capelin semen based on quantity. Completely stripped male capelin were kept in a tank, and every alternate day over a 6-day period, a subset of capelin was scored to assess their semen regeneration ability based on three quantity categories, each distinguished by a unique shade. The total number of capelin scored on specific dates is indicated above each respective bar. The y-axis represents proportionate scores, standardized through conversion to proportionate values, normalizing the data for comparison. The capture date and location are provided at the top.

**Fig. 7.**
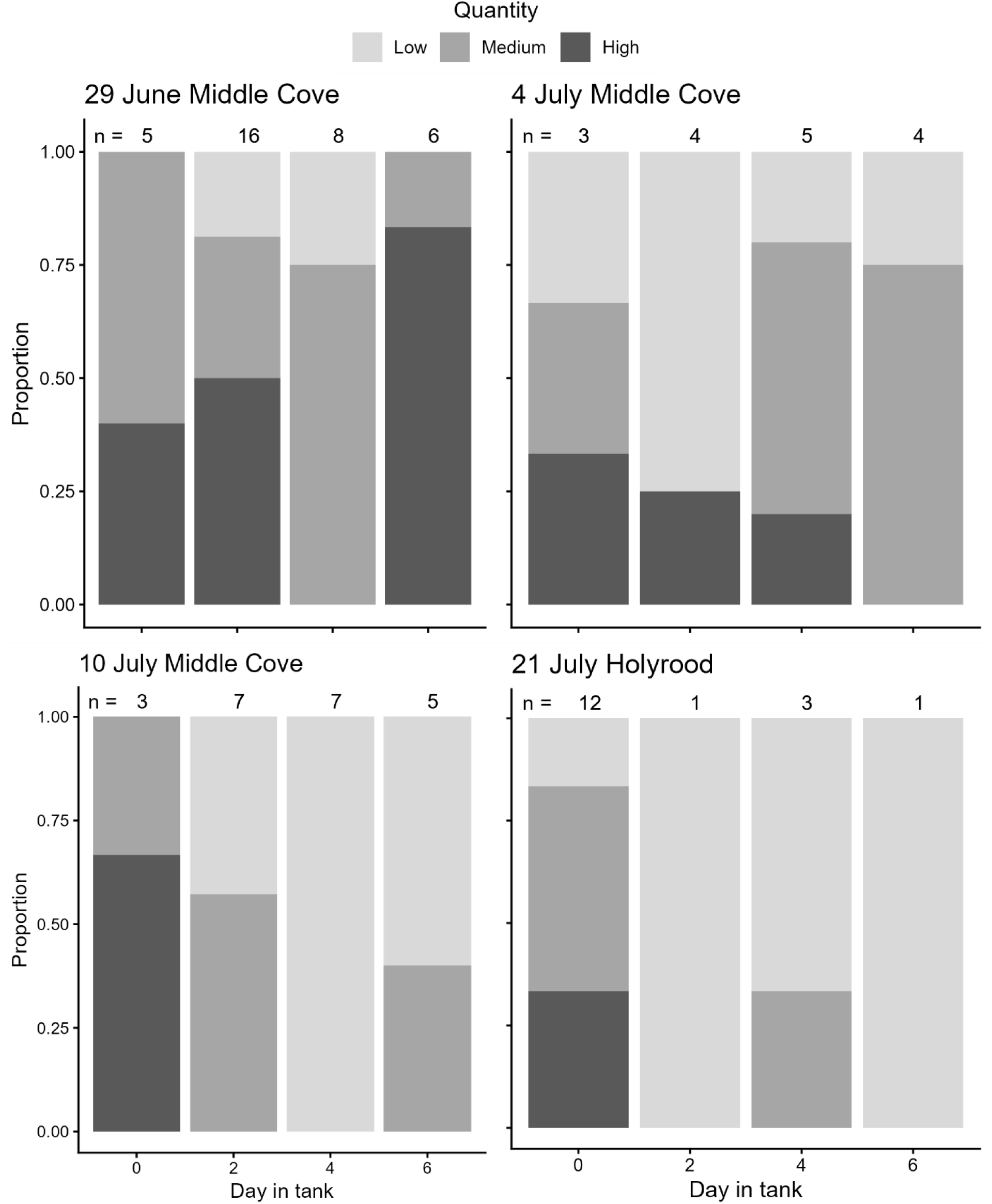
Assessment of the regenerative capacity of fair quality capelin semen based on quantity. Completely stripped male capelin were kept in a tank, and every alternate day over a 6-day period, a subset of capelin was scored to assess their semen regeneration ability based on three quantity categories, each distinguished by a unique shade. The total number of capelin scored on specific dates is indicated above each respective bar. The y-axis represents proportionate scores, standardized through conversion to proportionate values, normalizing the data for comparison. The capture date and location are provided at the top.

There was no relation between male size and the quality of regenerated semen (Fig. 8). Body size was not associated with the quantity of fair-quality or excellent quality regenerated semen (Fig. 9).

**Fig. 8.**
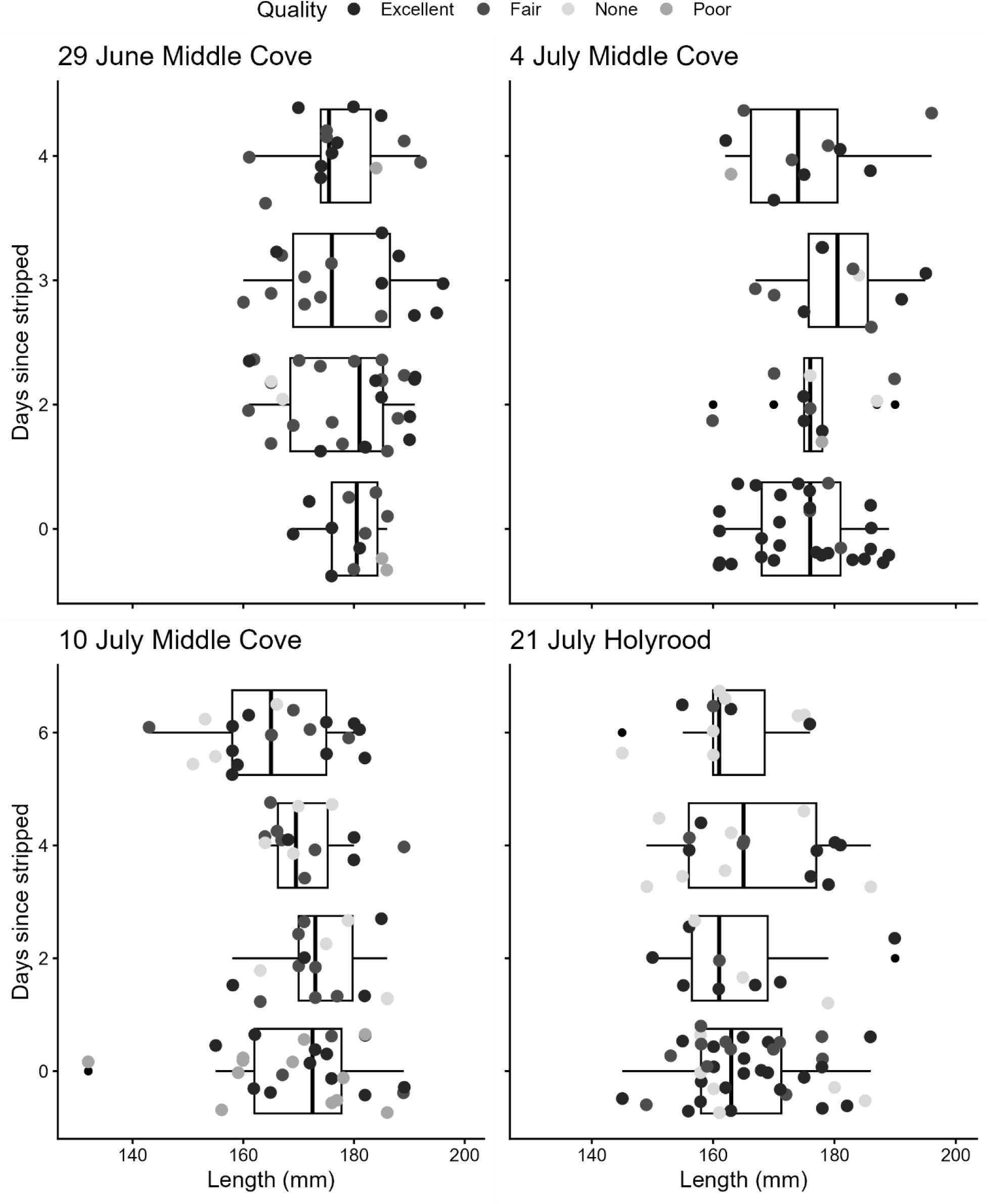
Variation in capelin semen regenerative capacity across different fish total lengths based on quality. Completely stripped capelin were kept in tanks, and a subset scored every alternate day over a 6-day period to assess semen regeneration. Semen quality is represented by size-differentiated shaded points, with larger and darker points denoting better quality. The thick line within the box plot represents the median, the box represents the interquartile range (25th and 75th quartiles), and the whiskers extend to 1.5 times the interquartile range from the first and third quartiles, respectively. The capture date and location are provided at the top.

**Fig. 9.**
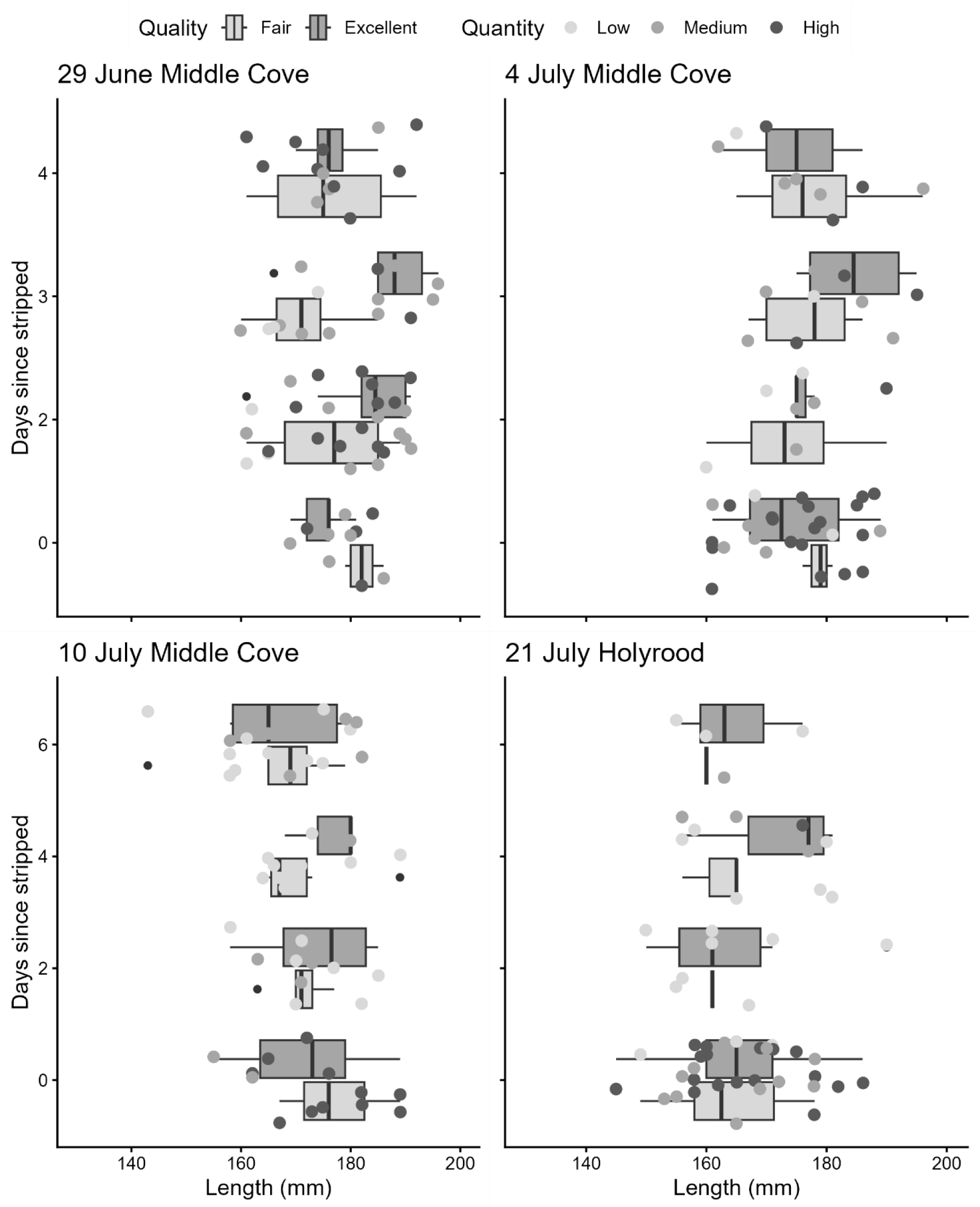
Variation in capelin semen regenerative capacity across different fish total lengths based on quantity. Completely stripped capelin were kept in tanks, and a subset scored every alternate day over a 6-day period to assess semen regeneration in the year 2023. Different box colours indicate semen quality, while semen quantity is represented by size-differentiated shaded points, with larger and darker points denoting better quantity. The thick line within the box plot represents the median, the box represents the interquartile range (25th and 75th quartiles), and the whiskers extend to 1.5 times the interquartile range from the first and third quartiles, respectively. The capture date and location are provided at the top.

## Discussion

Beach-spawning capelin males strategically position themselves near the beach, awaiting females to initiate spawning (Jeffers 1931; DFO 1991). Hence, males must be continuously prepared and should “compete” to extend their presence at spawning sites, a reproductive strategy known as endurance rivalry (Andersson 1994; Orbach et al. 2019). Our data on semen quality assessment in the field revealed that a majority of, but not all, capelin had adequate semen throughout the spawning season (not the same fish sampled). Furthermore, excellent semen quality was often accompanied by relatively high quantities. We found no relation between fish total length and semen quality, which is consistent with Orbach et al.’s (2020) findings that sperm swimming characteristics do not correlate with body length; however, the quantity of excellent quality semen was related to body size. As predicted, we found some capelin with no semen, but only towards the end of the sampling period. Previous work at the same study sites has, at times, observed many males that contain no semen (Purchase, unpublished). These results suggest that capelin progressively deplete their capacity to replenish semen throughout the spawning season.

Capelin—despite possessing unusually small testes (Orbach et al. 2020)—exhibited a strong capacity for semen regeneration, with the majority of captive fish regenerating semen within two days (we did not assess them after one day, except 29 June block). As different individuals were sampled each day, our design did not allow for the assessment of individual-level regeneration across multiple days, however, the trend suggests sampling on days 4 and 6 showed improved semen production compared to those sampled on day 2. Research on other teleosts such as brown trout (*Salmo trutta*), turbot (*Scophthalmus maximus*) and rainbow trout (*Oncorhynchus mykiss*), has shown that they also demonstrate semen regenerative capabilities, with sampling intervals set at biweekly for brown trout, and weekly for turbot and rainbow trout (did not investigate shorter intervals); however, upon repeated sampling of the same fish at each successive interval, these species experienced gradual declines in semen quality (Büyükhatipoglu and Holtz 1984; Suquet et al. 1992; Hajirezaee et al. 2009). Our data on greater semen regeneration on successive sampling may be due to our distinctive approach, where each fish was sampled only once for reassessment throughout the entire duration of captivity, meaning fish not sampled on day 2 had extra time for semen production. We found no relation between capelin body size and the quality and quantity of regenerated semen. However, we speculate that body size may influence how often semen regenerates (frequency of regeneration), warranting further investigation. Nevertheless, this rapid semen regeneration in capelin within two days improves mating opportunities and aligns with the concept of endurance rivalry, maximizing mating success.

Male-biased sexual size dimorphism in capelin is atypical compared to most teleosts as they lack physical combat during spawning. Thus, it may have evolved as an adaptation for endurance rivalry. Secondary sexual characteristics—enlarged pectoral, pelvic, and anal fins, and lateral ridges—are likely under natural selection by facilitating physical contact with females during mating rather than being under sexual selection (Orbach et al. 2019). Although, they could potentially aid in scramble competition if present. These traits are not mutually exclusive as larger body size allows males to persist at spawning sites, while secondary sexual characteristics contribute to mating success. Our results indicate that males possess the ability to rapidly regenerate semen, and thus, support the endurance rivalry hypothesis. Despite having unusually small testes and unique semen physiology (Beirão et al. 2015; Orbach et al. 2020), capelin maintain a high capacity for semen replenishment, allowing males to remain reproductively viable for longer periods, and thereby increasing the likelihood of additional mating opportunities.

## Acknowledgements

Assistance in collecting capelin and conducting experiments was provided by Kaitlyn Gladney. Funding was provided via Memorial University of Newfoundland and grants to Craig Purchase from the Natural Sciences and Engineering Research Council of Canada, the Canada Foundation for Innovation, and the Research and Development Corporation of Newfoundland and Labrador.

All procedures followed the Canadian Council on Animal Care guidelines for the use of research animals (Memorial University protocols 18-02-CP, 22-02-CP) and Fisheries and Oceans Canada permits (NL-6666-22 and NL-7297-23).

## References

Andersson, M. 1994. Sexual selection. Princeton University Press.

Beirão, J., Lewis, J.A., and Purchase, C.F. 2015. Spermatozoa ultrastructure of two osmerid fishes in the context of their family (Teleostei: Osmeriformes: Osmeridae). J. Appl. Ichthyol., 31(S1), 28–33. doi:10.1111/jai.12724

Beirão, J., Lewis, J.A., Wringe, B.F., and Purchase, C.F. 2018. A novel sperm adaptation to evolutionary constraints on reproduction: pre-ejaculatory sperm activation in the beach spawning capelin (Osmeridae). Ecol. Evol. 8(4): 2343–2349. doi:10.1002/ece3.3783.

Bisazza, A. 1993. Male competition, female mate choice and sexual size dimorphism in poeciliid fishes. Mar. Behav. Physiol. 23(1–4): 257–286. doi:10.1080/10236249309378869.

Brockmann, H. J. (2001). The evolution of alternative strategies and tactics (Vol. 30, pp. 1–51). Academic Press. doi:10.1016/S0065-3454(01)80004-8

Büyükhatipoglu, S., and Holtz, W. 1984. Sperm output in rainbow trout (Salmo gairdneri): effect of age, timing and frequency of stripping and presence of females. Aquaculture 37(1): 63–71. doi:10.1016/0044-8486(84)90044-9.

Castellano, S., Zanollo, V., Marconi, V., and Berto, G. 2009. The mechanisms of sexual selection in a lek-breeding anuran, Hyla intermedia. Anim. Behav. 77(1): 213–224. doi:10.1016/j.anbehav.2008.08.035.

Cox, R.M., Skelly, S.L., and John-Alder, H.B. 2003. A comparative test of adaptive hypotheses for sexual size dimorphism in lizards. Evolution 57(7): 1653–1669. doi:10.1111/j.0014-3820.2003.tb00371.x.

Darwin, C. 1872. The descent of man, and selection in relation to sex. Murray, London.

Dechaume-Moncharmont, F.-X., Brom, T., and Cézilly, F. 2016. Opportunity costs resulting from scramble competition within the choosy sex severely impair mate choosiness. Anim. Behav. 114: 249–260. doi:10.1016/j.anbehav.2016.02.019.

DFO. 1991. The science of capelin: a variable resource. Newfoundland Region. https://waves-vagues.dfo-mpo.gc.ca/Library/17629.pdf.

Dodson, J.J., Carscadden, J.E., Bernatchez, L., and Colombani, F. 1991. Relationship between spawning mode and phylogeographic structure in mitochondrial DNA of North Atlantic capelin Mallotus villosus. Mar. Ecol. Prog. Ser. 76(2): 103–113. doi:10.3354/meps076103.

Fairbairn, D.J. 1997. Allometry for sexual size dimorphism: pattern and process in the coevolution of body size in males and females. Annu. Rev. Ecol. Syst. 28(1): 659–687. doi:10.1146/annurev.ecolsys.28.1.659.

Hajirezaee, S., Amiri, B.M., and Mirvaghefi, A.R. 2009. Effects of stripping frequency on semen quality of endangered Caspian brown trout, Salmo trutta caspius. Am. J. Anim. Vet. Sci. 4(3): 65–71. doi:10.3844/ajavsp.2009.65.71.

Horne, C.R., Hirst, A.G., and Atkinson, D. 2020. Selection for increased male size predicts variation in sexual size dimorphism among fish species. Proc. R. Soc. B 287(1918): 20192640. doi:10.1098/rspb.2019.2640.

Jangaard, P.M. 1974. The capelin (Mallotus villosus): biology, distribution, exploitation, utilization, and composition. Bull. Fish. Res. Board Can. 186: 1–70.

Jeffers, G.W. 1931. The life history of the capelin Mallotus villosus (O.F. Müller). University of Toronto.

Orbach, D.N., Donovan, M., and Purchase, C.F. 2019. Sexually selected traits are larger and more variable in male than female beach-spawning capelin, Mallotus villosus. J. Fish Biol. 95(6): 1385–1390. doi:10.1111/jfb.14145.

Orbach, D.N., Rooke, A.C., Evans, J.P., Pitcher, T.E., and Purchase, C.F. 2020. Assessing the potential for post-ejaculatory female choice in a polyandrous beach-spawning fish. J. Evol. Biol. 33(4): 449–459. doi:10.1111/jeb.13579.

Parker, G. A. (1992). The evolution of sexual size dimorphism in fish. J. Fish Biol. 41(sB), 1–20. doi:10.1111/j.1095-8649.1992.tb03864.x

Penton, P.M., McFarlane, C.T., Spice, E.K., Docker, M.F., and Davoren, G.K. 2014. Lack of genetic divergence in capelin (Mallotus villosus) spawning at beach versus subtidal habitats. Can. J. Zool. 92(5): 377–382. doi:10.1139/cjz-2013-0261.

Pitt, T.K. 1958. Distribution, spawning and racial studies of the capelin, Mallotus villosus (Müller), in the offshore Newfoundland area. J. Fish. Res. Board Can. 15(3): 275–293.

Purchase, C.F., Evans, J.P., and Roncal, J. 2026. A framework for integrating natural and sexual selection within- and across-generations of eukaryotic biphasic life cycles. EcoEvoRxiv v3. doi:10.32942/osf.io/eu3am.

Pyron, M., Pitcher, T.E., and Jacquemin, S.J. 2013. Evolution of mating systems and sexual size dimorphism in North American cyprinids. Behav. Ecol. Sociobiol. 67(5): 747–756. doi:10.1007/s00265-013-1498-5.

R Core Team. 2023. *R: a language and environment for statistical computing*. R Foundation for Statistical Computing, Vienna, Austria. https://www.R-project.org/.

Rennie, M.D., Purchase, C.F., Lester, N., Collins, N.C., Shuter, B.J., and Abrams, P.A. 2008. Bioenergetic differences in energy acquisition and metabolism explain sexual size dimorphism in percids. J. Anim. Ecol. 77(5): 916–926. doi:10.1111/j.1365-2656.2008.01412.x.

Roff, D.A. 1986. Predicting body size with life history models: simple life history models can help evaluate the importance of ecological factors in the evolution of body size. BioScience 36(5): 316–323. doi:10.2307/1310236

Shine, R. 1988. The evolution of large body size in females: a critique of Darwin’s “fecundity advantage” model. Am. Nat. 131(1): 124–131.

Shuker, D.M., and Kvarnemo, C. 2021. The definition of sexual selection. Behav. Ecol. 32(5): 781–794. doi:10.1093/beheco/arab055.

Shine, R., Olsson, M.M., Moore, I.T., LeMaster, M.P., Greene, M., and Mason, R.T. 2000. Body size enhances mating success in male garter snakes. Anim. Behav. 59(3): F4–F11. doi:10.1006/anbe.1999.1338.

Shine, R., Langkilde, T., and Mason, R.T. 2003. Confusion within mating balls of garter snakes: does misdirected courtship impose selection on male tactics? Anim. Behav. 66(6): 1011–1017. doi:10.1006/anbe.2003.2301.

Suquet, M., Omnes, M.H., Normant, Y., and Fauvel, C. 1992. Influence of photoperiod, frequency of stripping and presence of females on sperm output in turbot, Scophthalmus maximus. Aquac. Res. 23(2): 217–225. doi:10.1111/j.1365-2109.1992.tb00612.x.

Templeman, W. 1948. The life history of the capelin (Mallotus villosus O.F. Müller) in Newfoundland waters. Bull. Nfld. Gov. Lab. 17: 1–151.

Trivers, R.L. 1972. Parental investment and sexual selection. In Sexual selection and the descent of man 1871–1971. Edited by B. Campbell. Aldine, Chicago. pp. 136–179.

Wagle, R. 2024. Pre- and post-mating selection on male capelin. M.Sc. thesis, Memorial University of Newfoundland.

Webb, T.J., and Freckleton, R.P. 2007. Only half right: species with female-biased sexual size dimorphism consistently break Rensch’s rule. PLoS ONE 2(9): e897. doi:10.1371/journal.pone.0000897.

Winters, G.H. 1970. Biological changes in coastal capelin from overwintering to spawning condition. J. Fish. Res. Board Can. 27(12): 2215–2224.

